# The latitudinal gradient in rates of evolution for bird beaks, a species interaction trait

**DOI:** 10.1101/2020.07.31.231142

**Authors:** Benjamin G Freeman, Dolph Schluter, Joseph A Tobias

## Abstract

Where is evolution fastest? The biotic interactions hypothesis proposes that greater species richness creates more ecological opportunity, driving faster evolution at low latitudes, whereas the “empty niches” hypothesis proposes that ecological opportunity is greater where diversity is low, spurring faster evolution at high latitudes. Here we tested these contrasting predictions by analyzing rates of bird beak evolution for a global dataset of 1141 sister pairs of birds. Beak size evolves at similar rates across latitudes, while beak shape evolves faster in the temperate zone, consistent with the empty niches hypothesis. We show in a meta-analysis that trait evolution and recent speciation rates are faster in the temperate zone, while rates of molecular evolution are slightly faster in the tropics. Our results suggest that drivers of evolutionary diversification are more potent at higher latitudes, thus calling into question multiple hypotheses invoking faster tropical evolution to explain the latitudinal diversity gradient.

## Introduction

The biotic interactions hypothesis proposes that strong interactions between tropical species are both a cause and a consequence of the latitudinal diversity gradient (Schemske 2009). Imagine that diversity was equal in the tropics and temperate zone. If so, then stronger species interactions in the tropics are proposed to lead to faster adaptation and speciation at low latitudes, generating an excess of tropical species. This is because species interactions are thought to be the dominant drivers of selection for tropical organisms, and adaptation to species interactions is expected to lead to faster adaptive evolutionary change than adaptation to abiotic challenges (Dobzhansky 1950; MacArthur 1972; Schemske 2009). Now imagine that the latitudinal diversity gradient already exists. A greater number of species in the tropics could provide increased ecological opportunity via an increase in the diversity of possible interactions between species and greater spatial heterogeneity in these interactions, also leading to faster evolution (Schemske 2009). In this conception, diversity begets diversity in a positive feedback loop. Whether species interactions are stronger in the tropics has been tested by many studies, with mixed results (Schemske *et al*. 2009; Anstett *et al*. 2016; Moles & Ollerton 2016; Roslin *et al*. 2017; Baskett & Schemske 2018; Hargreaves *et al*. 2019; Freeman *et al*. 2020; Roesti *et al*. 2020). However, the prediction that stronger species interactions or higher diversity can drive faster evolution in the tropics remains largely unexplored (but see Benkman 2013).

An alternative hypothesis predicts that evolution is instead fastest where species diversity is low. According to standard niche theory, ecological opportunity should be negatively related to diversity (Hutchinson 1959), such that evolutionary rates are faster in regions with the fewest species (Lawson & Weir 2014; Schluter 2016; Schluter & Pennell 2017; Machac 2020). A geographic prediction arising from this “empty niches” hypothesis is that evolutionary rates should be faster in the species-poor temperate zone than the species-rich tropics. Glacial cycles are a prominent explanation for why species richness is currently low in the temperate zone despite its large area (Dynesius & Jansson 2000), with abundant empty niche space likely available following the periodic retreat of ice-sheets (Schluter 2016).

Here, we investigate the contrasting predictions of the biotic interactions and empty niche hypotheses by analyzing latitudinal differences in rates of bird beak evolution. Bird beak morphology can be linked to abiotic factors such as temperature because beak size functions in thermoregulation (Tattersall *et al*. 2009, 2017). However, beak morphological form—particularly shape—is tightly coupled with diet across all ~10,000 species of birds (Pigot *et al*. 2020). Indeed, beak shape predicts dietary niches even when they are subdivided into finely grained categories, suggesting a pervasive role for trophic adaptations as drivers of bird beak evolution (Pigot *et al*. 2020). These dietary functions mean the avian beak mediates species interactions both within and between trophic levels. On one hand, birds use their beaks to feed on prey animals and food plants, with beak morphology intimately shaped by these trophic interactions (Smith 1987; Benkman 2003; Lerner *et al*. 2011; Grant & Grant 2014; Cattau *et al*. 2018). On the other hand, competitive interactions between closely related species may also drive beak morphological evolution via ecological character displacement (Brown & Wilson 1956; Schluter 2000; Grant & Grant 2006). The biotic interactions hypothesis predicts that both types of interaction—trophic and competitive—accelerate beak evolution in the tropics.

We tested whether empirical patterns of bird beak evolution better matched predictions of the biotic interactions or empty niche hypotheses by compiling new measurements of beak size and shape (Tobias et al. this volume) for a previously published global dataset of bird sister pairs (Cooney *et al*. 2017b). First, we tested whether beak size and shape evolution are fastest for tropical sister pairs, as expected under the biotic interactions hypothesis, or fastest for temperate sister pairs, as expected under the empty niches hypothesis. Second, we isolated the effect of competitive interactions on latitudinal differences in beak evolution by examining rates of beak evolution in sympatric sister pairs. Interspecific competition can lead to greater trait divergence in sympatry than in allopatry via two processes: ecological character displacement, where competition directly drives trait divergence (Stuart & Losos 2013; Weber & Strauss 2016), and ecological sorting, where competition prevents sympatry between species with similar traits but permits sympatry between species with divergent traits (Pigot & Tobias 2013; Tobias *et al*. 2014, 2020; Freeman 2015). The prediction of the biotic interactions hypothesis is that competitive interactions resulting in ecological character displacement or ecological sorting are strongest in the tropics. If so, then evolutionary rates for sympatric sister pairs, relative to baseline rates for allopatric sister pairs, should be faster in the tropics than in the temperate zone. The empty niches hypothesis makes no clear prediction about latitudinal gradients in the strength of competition between close relatives.

Our analysis of global patterns of bird beak evolution sheds light on the more general problem of why species richness is highest in the tropics. Dozens of hypotheses have been proposed to explain the latitudinal diversity gradient. The biotic interactions hypothesis is one of at least seven distinct hypotheses that invoke faster tropical evolution to explain why species richness peaks near the equator (Mittelbach *et al*. 2007). These “fast tropical evolution” hypotheses propose different mechanisms for why evolution is most rapid in the tropics, including stronger species interactions (a focus of this manuscript), faster rates of molecular evolution (Rohde 1992; Gillman & Wright 2014), narrower physiological tolerances (Janzen 1967), larger spatial area (Rosenzweig 1995), greater climatic variation (Haffer 1969), greater genetic drift (Fedorov 1966), and greater divergence in the face of gene flow (Moritz *et al*. 2000). Finding faster rates of beak evolution in the tropics would therefore support the biotic interactions hypothesis while also being consistent with the entire set of fast tropical evolution hypotheses. A result that beak evolution is faster in the temperate zone, however, would support the empty niches hypothesis while calling into question all hypotheses based on the concept of faster tropical evolution.

## Materials and methods

### Sister pair selection and geographic variables

We used a previously published dataset of 1306 sister pairs of birds with information on median latitude and range overlap for each pair (Cooney *et al*. 2017b). This dataset defined sister pairs from a dated global bird phylogeny of species, subsetted to include only species placed on the tree on the basis of genetic data (6670 species out of 9993) (Jetz *et al*. 2012). Each sister pair in this dataset comprises two species that are each other’s closest relatives. Because of incomplete sampling, not all sister pairs are true sister species. Nevertheless, each sister pair is a phylogenetically independent comparison between two closely related species, and is therefore appropriate for our purposes. Because our primary goal is to compare rates of beak evolution in tropical versus temperate sister pairs, we excluded 141 sister pairs from mixed latitudinal zones, defined as cases where one species was classified as tropical (midpoint latitude < 23.4°) and the other as temperate (midpoint latitude > 23.4°). We further excluded 14 sister pairs with estimated divergences greater than 20 million years, and 10 sister pairs where morphological data was lacking for one species within the pair. Our final dataset contained 1141 sister pairs.

Three key variables in this dataset are: (1) divergence time; (2) latitude; and (3) range overlap, coded as allopatric vs. sympatric. We used divergence time—the time to most recent common ancestor estimated using the dated phylogeny—as an estimate of evolutionary age. For latitude, we calculated the median latitude for each species in our dataset based on standard geographical range polygons from BirdLife International (BirdLife International 2018). Following numerous previous studies (e.g., Weir & Schluter 2007; Cooney *et al*. 2017b; Sheard *et al*. 2020), we focused solely on the breeding distribution, where interactions over food and territories are likely most intense, and averaged the latitude for each pair of sister species to produce a midpoint latitude for the pair. Using these midpoint latitudes, we then classified each sister pair as tropical (< 23.4°; N = 800) or temperate (> 23.4°; N = 341).

Range overlap is a continuous variable ranging from 0, when the two species in the pair do not overlap in range, to 1, where the range of the smaller-ranged species is completely subsumed within the range of the larger-ranged species. Following previous studies (e.g., Tobias *et al*. 2014; Cooney *et al*. 2017b), we used a threshold of 20% range overlap to define whether sister pairs were allopatric (< 20% range overlap; N = 746) or sympatric (> 20% range overlap; N = 395). Hence, sister pairs were coded as sympatric only when they had substantial range overlap such that the two species likely interact. A range overlap of 20% often spans a large geographic area. In continental systems, any interaction-driven trait divergence within this region of overlap may extend far into the allopatric zone because of unimpeded gene flow (e.g., Kirschel *et al*. 2019). To examine the sensitivity of results to a 20% overlap threshold, we conducted further tests using thresholds of > 0% and > 50% range overlap to define sympatry and allopatry. Finally, we note that beak measurements were sampled explicitly in the area of range overlap for some sister pairs (e.g., Furnariidae; Tobias *et al*. 2014) but this was not possible for all pairs. Further studies sampling morphology in both sympatric and allopatric populations for each sister pair would be necessary to directly test patterns of ecological character displacement.

### Morphological divergence

We extracted morphological data for each species in each species pair from a comprehensive global dataset of bird morphology (Tobias et al. this volume). We analyzed four traits, all measured in mm: (1) beak length along the culmen from the skull to the tip; (2) beak length from the nares to the tip; (3) beak depth at the nares; and (4) beak width at the nares. We excluded individuals that had missing values for any beak traits. Our final morphological dataset contained 9,966 individuals (mean 4.4 individuals per species; range = 1–89; 94% of species had four or more individuals measured).

We used a principal components analysis to place all individuals within a single multidimensional beak morphospace. To generate independent axes of variation in beak morphology, we ran a principal component analysis (PCA) on logged values of each of the four beak traits. The resulting morphospace is essentially two dimensional: PC1, related to overall beak size, explained 82.1% of variation, while PC2, a metric of beak shape, explained a further 14.3% of variation (Table S1, Figure S1). We quantified the volume of beak morphospace for tropical and temperate zone species by constructing a hypervolume using Gaussian kernel density estimation and calculating its volume using the “hypervolume” package (Blonder *et al*. 2014; Blonder & Harris 2019). Tropical species outnumber temperate species by approximately 3 to 1 in this dataset, reflecting the overall global ratio. However, the volumes of beak morphospace occupied by tropical and temperate species in this dataset are similar (four-dimensional volumes of 4.55 versus 4.91; see Figure S2 for a two-dimensional representation). This implies that tropical species are more densely packed within beak morphospace than temperate species, a feature found in other avian diversity gradients (Pigot *et al*. 2016). We calculated divergence in beak size (PC1) and shape (PC2) for each taxon pair as the absolute value of the difference in species means along these axes. Sampling error will cause the raw difference in species’ means to be larger on average than the true difference in species means. We therefore estimated bias-corrected divergence in beak size and shape for each sister pair using pooled sample variance calculated for all species. For demonstration of this bias, as well as code to calculate bias-corrected divergence, see R scripts provided in the Supporting Information. Following bias correction, some values of beak size and shape divergence between sister pairs are negative.

### Statistical analysis

We tested the hypothesis that evolutionary rates of beak divergence differ between latitudes by fitting evolutionary and empirical models to our trait data in R version 3.9 (R Development Core Team 2020). We fit distinct models to divergence in beak size (PC1) and divergence in beak shape (PC2). For all beak size models, we removed two outlier sister pairs (the temperate pair *Numenius minutus* and *N. arquata* (curlews) and the tropical pair *Geospiza magnirostris* and *G. fuliginosa* (Darwin’s finches from the Galápagos) with extremely high beak size divergence but very low estimated divergence dates. These cases likely arise from phylogenetic error.

We first fit Brownian motion (BM) and Ornstein Uhlenbeck (OU) models to bias-corrected morphological differences and time. These common models of trait evolution pass through the origin. In our dataset, however, both beak size and beak shape divergence are relatively high even at very young evolutionary ages (Figure 1a and 1b), as shown in previous sister species analyses (McEntee *et al*. 2018). There are three possible explanations for this seemingly impossible initial rate of evolution. The first is statistical: evolutionary age is estimated with error, and any species pairs whose ages are underestimated will lead to models with an intercept fitting better than models fit through the origin. The second explanation is biological: rapid evolution and phenotypic plasticity can lead to trait differences that arise prior to divergence at the putatively neutral loci that are used to estimate evolutionary age. Third, it is possible that sister pairs with high morphological divergence but very young ages represent cases where sister species are moderately old but recent introgression has created the illusion of evolutionary youth (Tobias *et al*. 2020).

**Figure 1.**
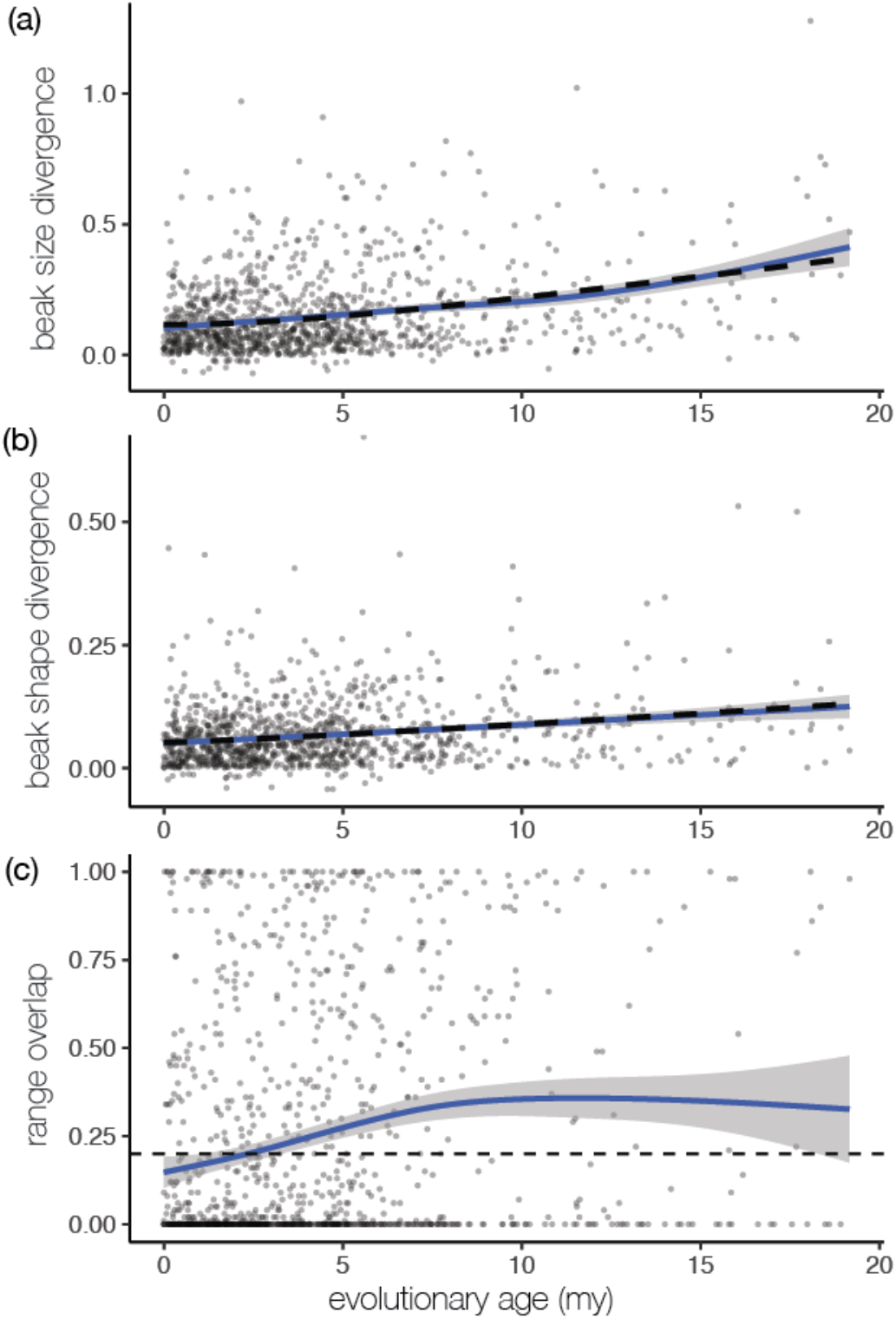
Patterns of divergence in beak size (a), beak shape (b) and range overlap (c) in 1141 sister pairs of birds. Raw data are plotted. Loess regressions are shown in blue; predictions from the best-fit models (power functions with intercepts) are shown as dashed black lines. Beak size and shape divergence values are corrected for bias arising from sampling error. Range overlap is the proportion of the smaller-ranged species that falls within the range of the larger-ranged species. Sister pairs were coded as sympatric if they had range overlaps > 0.20 (the dashed line).

We therefore explored fitting simple models with intercepts that better captured the patterns in our data than formal evolutionary models. Because a high intercept is consistent with a high rate of evolution in the early stages of divergence (McEntee *et al*. 2018), we compare intercepts as well as slopes of fitted models. We included simple power functions in the set of models because they are more flexible than the evolutionary models, and can fit relationships suggested by the loess regressions shown in Figure 1. In sum, we fit five models: BM and OU models, a BM model with an intercept, a power function forced through the origin, and a power function with an intercept. We compared model fit using AIC.

The response variable in our models (either beak size divergence or beak shape divergence) was the signed square root of squared differences between bias-corrected species means divided by two. BM and OU models estimate the variance between species times two on the squared differences scale; we correct for this by dividing by two. We used the signed square root of squared differences—the visual scale for analyzing trait differences—because residuals are more normally distributed and had more homogenous variance on this scale. All results from models fit to squared differences data, weighting by the variance of residuals, were similar, but the skewed distributions of residuals on the squared difference scale led us to be cautious in interpretation of AIC scores and p-values (Figures S3 and S4, Tables S2 and S3).

We tested whether rates of trait evolution differed between the tropics and temperate zone by fitting models with and without a latitudinal zone term (temperate or tropical), and conducting *F* tests using the “anova” command in R. For the BM intercept and power function with intercept, we fit three models in which (1) neither the intercept nor the slope varied between tropical and temperate zone sister pairs; (2) the intercept varied between tropical and temperate zone sister pairs; and (3) both slope and intercept varied between tropical and temperate zone sister pairs.

Next, we analyzed whether sympatry is associated with greater rates of beak size and beak shape evolution in our dataset (Figure 1c). We began with the best-fit model from our previous analysis (power function with intercept). We then fit two additional models in which (1) the intercept varied between allopatric and sympatric sister pairs; and (2) both slope and intercept varied between allopatric and sympatric sister pairs. We compared these models by conducting *F* tests using the “anova” command in R.

The key prediction arising from the biotic interactions hypothesis is that evolutionary rates in sympatry are particularly fast in the tropics. To test this prediction, we added a latitudinal zone term to the best-fit model that included a sympatric/allopatric term, as described in the previous paragraph. We tested models in which (1) the intercept varied between tropical and temperate zone sister pairs; and (2) both slope and intercept varied between tropical and temperate zone sister pairs. Again, we compared model fit by conducting *F* tests using the “anova” command in R.

## Literature review

To help place the results of our study in a broader context, we conducted a meta-analysis of previous studies of latitudinal gradients in evolutionary rates. BGF conducted Web of Science and Google Scholar searches on 5 November 2020 for “molecular evolution AND latitud*, “trait evolution AND latitud*”, “speciation AND latitud*”, and “evolution AND latitud*”, and retained studies that estimated evolutionary rates in both the tropics and temperate zone. We categorized studies as measuring three types of evolutionary rates: rates of molecular evolution, rates of trait evolution, or recent speciation rates. Theory has shown that speciation rates inferred from lineage-through-time plots may be unreliable (Louca & Pennell 2020) but that speciation rates close to the present can be unambiguously identified because there is no confounding effect of extinction. We therefore included only studies that estimated recent speciation rates for both the tropics and temperate zone. For each study, we calculated the ratio in evolutionary rates in the temperate zone versus evolutionary rates in the tropics (details provided in Supporting Information). We were unable to calculate standard errors for the majority of studies and therefore could not calculate fit a random effects meta-analysis to the data. Instead, we calculated weighted averages for the ratio of temperate versus tropical rates for each type of evolutionary rate.

## Results

We found evolutionary rates were faster in the temperate zone for beak shape and similar between tropical and temperate zone birds for beak size (Figures 2 and S5). For both beak size and beak shape, the best-fit model was a power function with an intercept, and standard evolutionary models (Brownian motion and Ornstein Uhlenbeck) were poor fits (Δ AIC > 40; Table 1). For beak shape but not beak size, models that included a latitudinal zone term (tropical/temperate) improved model fit (e.g., for the best-fit power function with intercept, *p* = 0.0034; Table S4), with higher intercepts for beak shape divergence in the temperate zone. We found further evidence that the slope of beak shape divergence is also steeper in the temperate zone (*p* = 0.039; model where both intercept and slope differ between tropics and temperate zone plotted in Figure 2b). Young lineages in the temperate zone have greater beak shape divergence than equivalently aged lineages in the tropics (higher intercept), but successively older lineages in the temperate zone have even greater beak shape divergence than equivalently aged lineages in the tropics (steeper slope).

**Figure 2.**
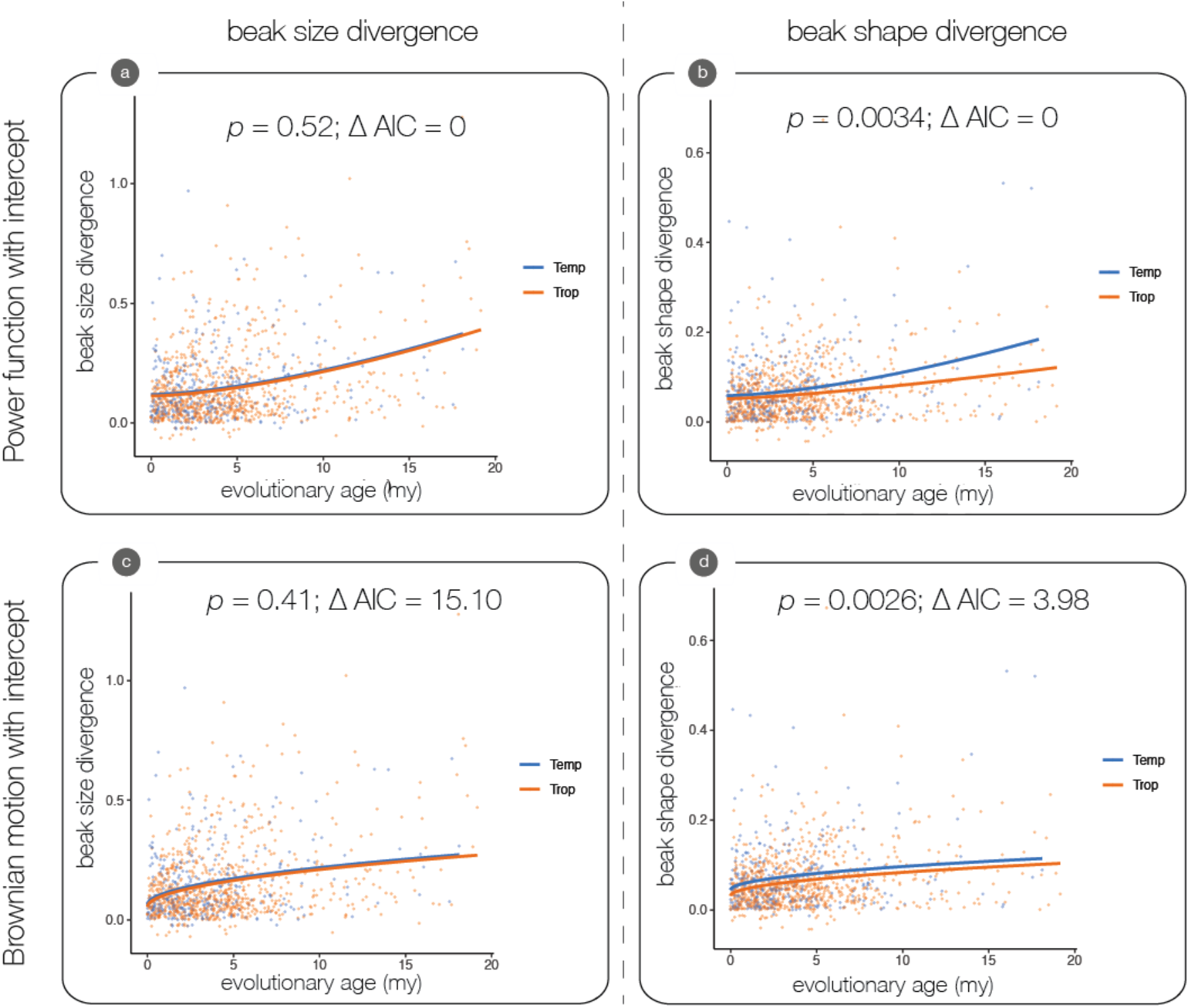
Rates of beak size evolution are similar between tropics and temperate zone (left panels), but beak shape evolution is faster in the temperate zone (right panels). Model predictions are plotted for the two best models: power functions with an intercept (a, b) and Brownian motion models with an intercept (c, d). *P*-values are from *F* tests testing whether the inclusion of a tropical/temperate term improved model fit. Δ AIC values compare different model fits separately for beak size and beak shape.

**Table 1.**
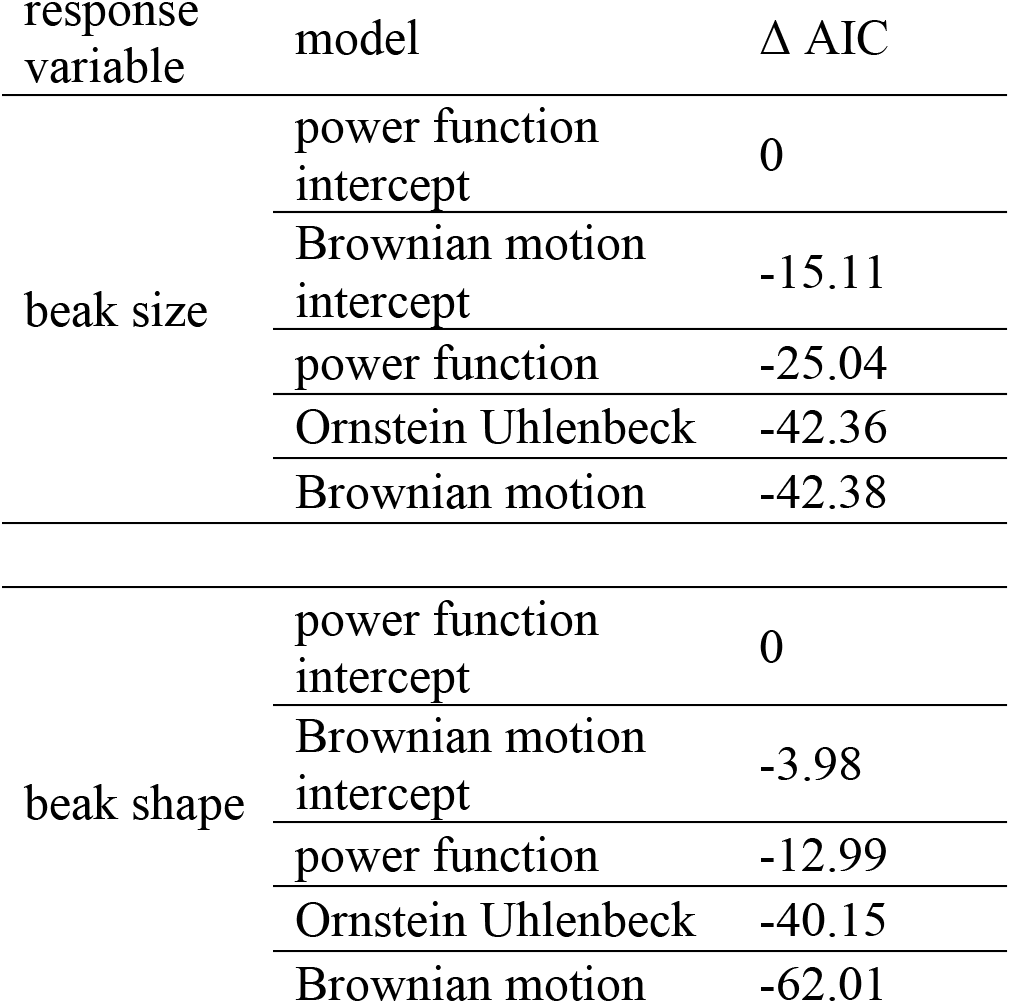
Models that included intercepts (power function with an intercept and Brownian motion model with an intercept) were better fits than models forced through the origin (power function, Ornstein Uhlenbeck and Brownian motion models).

Sympatric sister pairs have greater trait divergence than allopatric sister pairs for beak size (Figure 3a, t-test *p* = 0.020, means = 0.17 and 0.14, respectively) and beak shape (Figure 3b, t-test *p* < 0.001, means = 0.077 and 0.061, respectively). This pattern of greater divergence in sympatry is consistent with a role for competition in driving trait divergence. However, beak size and beak shape divergence also increase with evolutionary age (Figure 1 a,b), and sympatric sister pairs are older than allopatric sister pairs. In this dataset, median sister pair ages are 2.90 and 4.54 million years for tropical allopatric and sympatric sister pairs, respectively (t-test on mean age; *p* < 0.001), and 2.30 and 3.60 million years for temperate zone allopatric and sympatric sister pairs (t-test on mean age; *p* = 0.0039; Figure 3).

**Figure 3.**
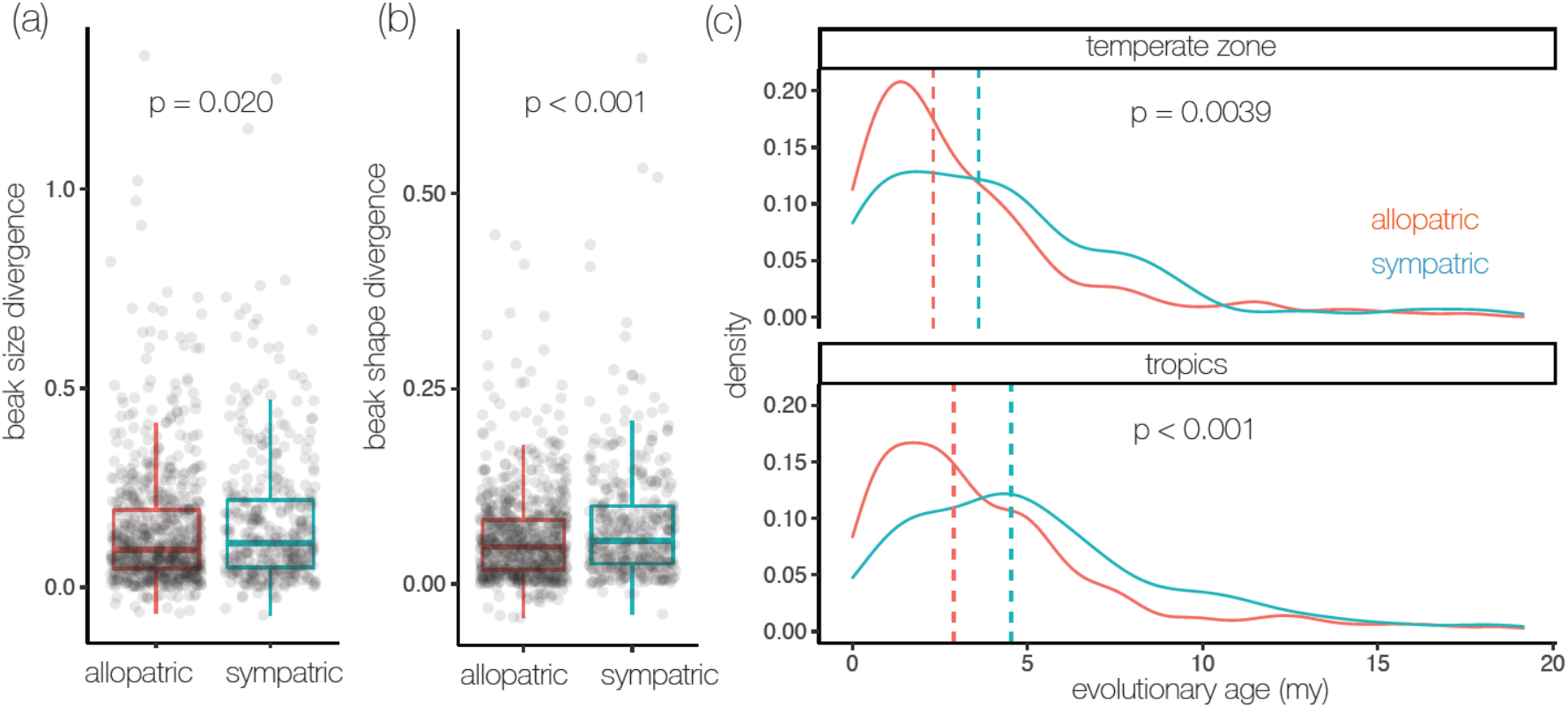
Sympatric sister pairs have greater beak size divergence (a), beak shape divergence (b), and are older (c) than allopatric sister pairs. *P*-values are from t-tests of trait divergence or ages between allopatric and sympatric sister pairs; separate t-tests for age for temperate and tropical zones in panel (c). Median values of ages for temperate and tropical sister pairs are plotted as vertical dashed lines.

Statistically controlling for evolutionary age, we found evidence that competition drives beak shape evolution, but not beak size evolution. For beak shape, including a range overlap term (sympatry/allopatry) improved model fit (*p* = 0.0022, Figure 4b, Table S5). In contrast, models that included a range overlap term did not improve model fit for beak size (*p* = 0.20, Figure 4a, Table S5). This indicates that the greater divergence in beak size for sympatric sister pairs is primarily due to their older age. Models that included a temperate/tropical term in addition to a sympatry/allopatry term further improved fit for beak shape (*p* = 0.0032, Figure 4d) but not for beak size (*p* = 0.69, Figure 4c, see Table S5). However, excess rates of divergence in sympatry compared to allopatry for beak shape were similar between the tropics and temperate zone (Figure 4d). Results were similar when using alternative thresholds to define sympatric versus allopatric sister pairs (Figures S6 and S7, Tables S6 and S7).

**Figure 4.**
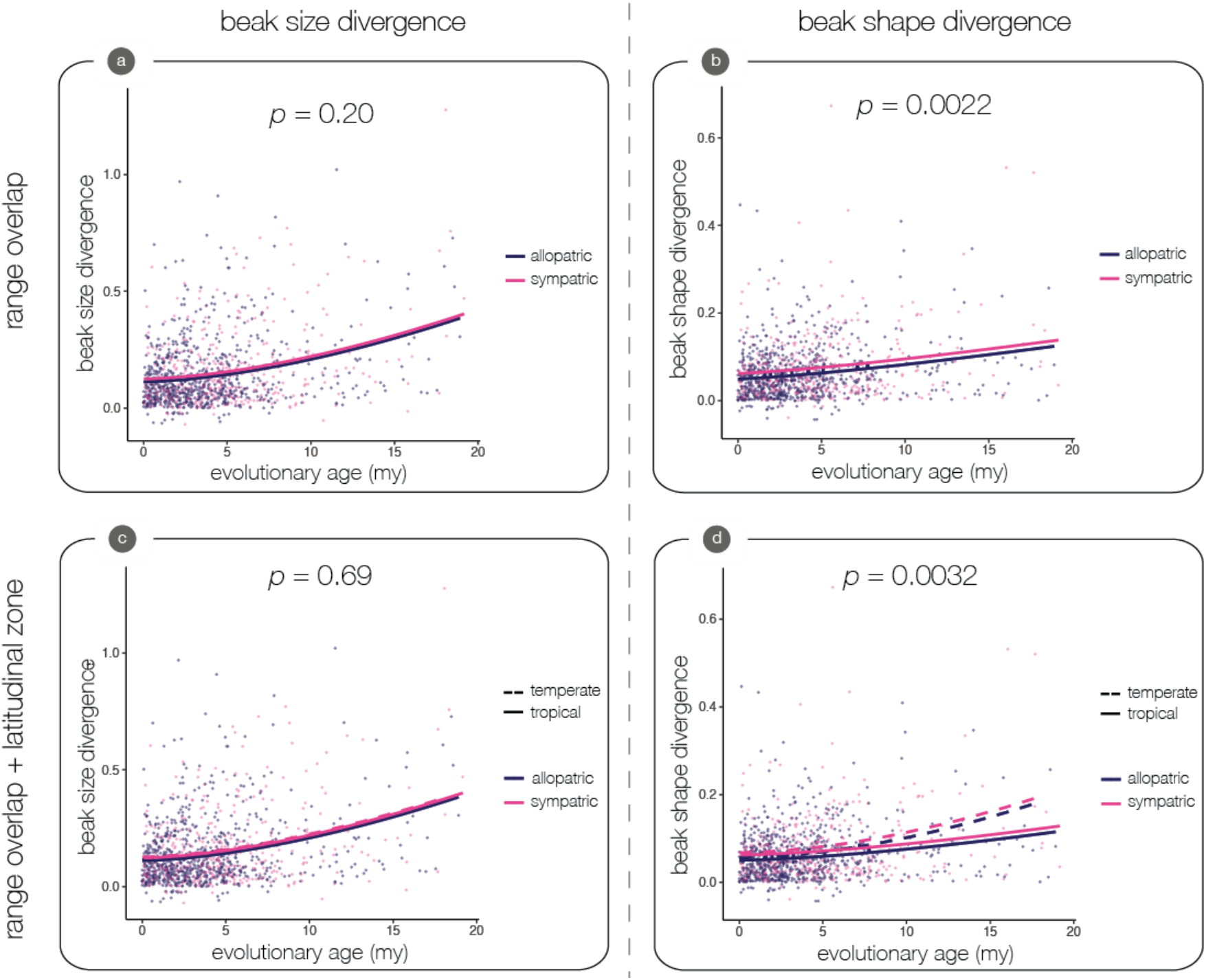
Rates of beak size evolution are similar between allopatric (n = 746) and sympatric (n = 395) sister pairs (left panels), but beak shape evolution is faster in sympatry (right panels). *P*-values are from *F* tests testing whether the inclusion of an allopatric/sympatric term to a power function with an intercept improved model fit (a, b), or whether the inclusion of a tropical/temperate term to a power function with an intercept and an allopatric/sympatric term improved model fit (c, d). The *p*-value for beak shape (d) is from a *F* test comparing a reduced model with an allopatric/sympatric term to a full model with terms allowing both the intercept and slope to differ between tropics and temperate zone. Beak shape evolution is faster in the temperate zone in both allopatry and sympatry compared to the tropics.

Our survey of published literature shows that the average ratio of evolutionary rates in the temperate zone versus the tropics is 1.46 ± 0.16 (mean ± standard error, weighted by sample size; N = 18 studies). This means that evolution is typically fastest in the temperate zone (Figure 5). Trait evolution and recent speciation rates are faster in the temperate zone (weighted means of 2.57 ± 1.45 and 1.50 ± 0.31, respectively), while rates of molecular evolution are slightly faster in the tropics (weighted mean of 0.92 ± 0.02).

**Figure 5.**
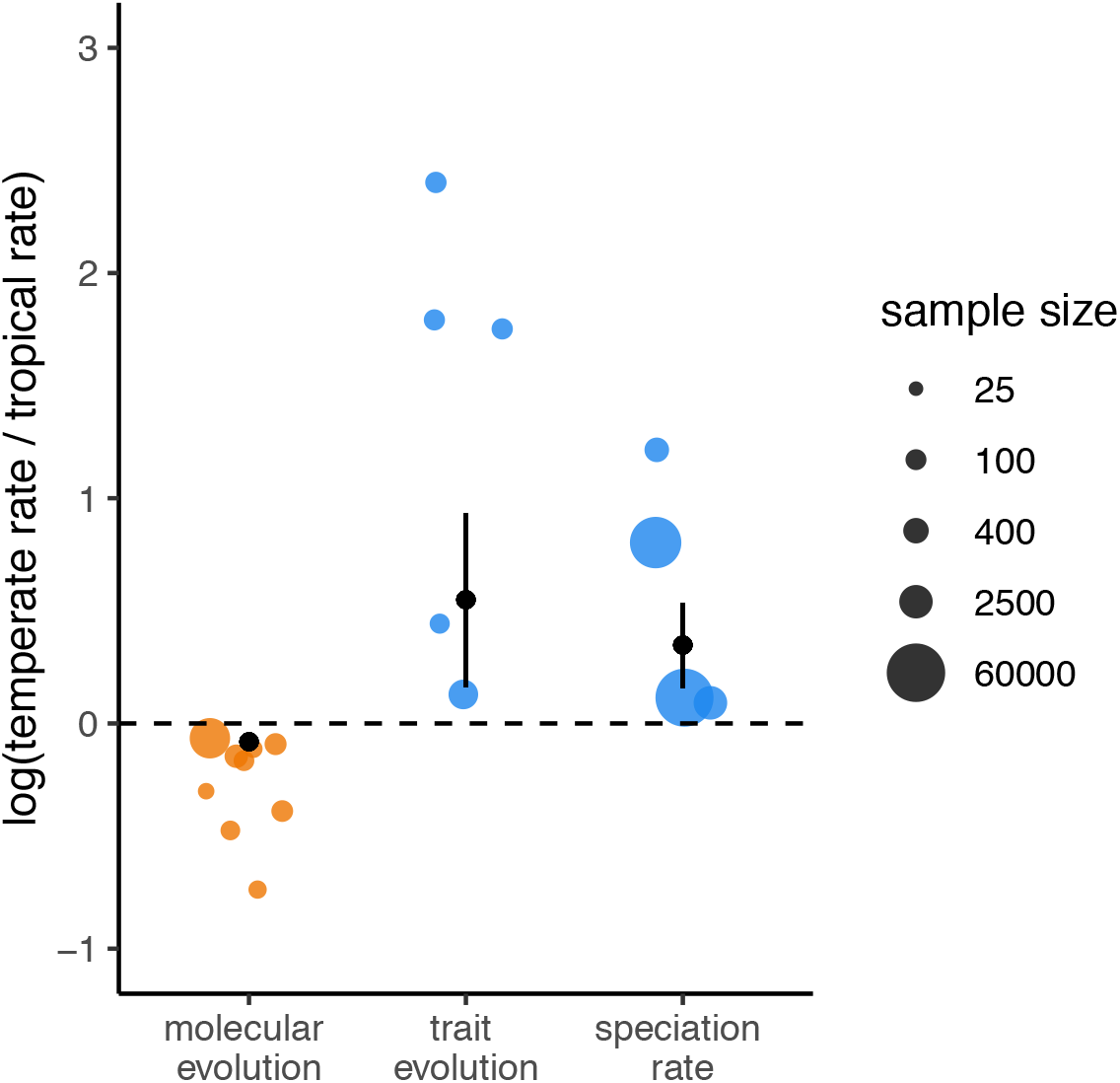
Meta-analysis of 18 published studies estimating latitudinal differences in evolutionary rates. Trait evolution and recent speciation rates are faster in the temperate zone, while rates of molecular evolution are slightly faster in the tropics. The weighted average and standard error for each category of evolutionary rate are illustrated as black points and lines. Point estimates of the ratio of temperate zone rate and tropical rate for each study are illustrated in blue when evolutionary rates are faster in the temperate zone and orange when faster in the tropics; size of points reflects sample size. We plot logged values to illustrate fold-differences between evolutionary rates from the temperate zone and tropics. See Supporting Information for details on calculating ratios for each study.

## Discussion

Our analyses of evolutionary rates across 1141 pairs of sister species reveal that bird beak shape evolution is fastest in the temperate zone, whereas bird beak size evolves at a similar rate across latitudes. In addition, sympatric tropical sister pairs do not have particularly fast evolutionary rates, as predicted if one specific biotic interaction—competition between closely related species that leads to exaggerated trait differences in sympatry—were stronger in the species-rich tropics. These results contradict both aspects of the biotic interaction hypothesis: that stronger biotic interactions in the tropics lead to faster evolution in biotic interaction traits, and that greater species richness leads to faster evolution in the tropics. Diversity does not beget diversity, at least for beak morphology. Instead, our results are more consistent with the empty niches hypothesis, whereby ecological opportunity promotes greater beak divergence in the species-poor temperate zone.

It is perhaps unsurprising that bird beak evolution is not fastest in the tropics where species richness is highest. Previous studies have shown that greater species richness is primarily associated with greater niche packing within beak morphospace rather than with niche expansion (Pigot *et al*. 2016). Indeed, beak morphospace is only slightly larger in the tropics in our dataset despite three times more species in the tropics. This increased saturation of available niche space should offer fewer opportunities for rapid beak evolution (Tobias *et al*. 2020). In addition, bird beaks are well described by a small number of dimensions and tend to reflect rather diffuse (one-to-many) interactions between consumers and a wide range of food plants or prey species. Diversity may beget diversity in other situations, however. For example, greater diversity could spur rapid evolution (and coevolution) in species interaction traits shaped by more specialized (e.g. one-to-one) interactions, such as the foraging traits of herbivorous beetles adapted to host trees (McKenna *et al*. 2009) or greater dimensionality, such as venoms that predators use to capture prey (Daltry *et al*. 1996) or cocktails of chemicals that plants use to deter herbivores (Rasmann & Agrawal 2011). Further studies that measure rates of evolution along the latitudinal gradient for traits that differ in their specialization or dimensionality would test these ideas.

A further important caveat relates to the precision of our trait quantification. Our data are based on a set of linear beak measurements and thus provide only a partial estimate of aspects such as curvature. The beaks of two interacting hummingbird species may potentially diverge in the extent of curvature linked to different floral niches while appearing identical from the perspective of our measurements. It is clear that the linear measurements we use nonetheless explain most of the ecological variation among species, including relatively subtle behavioral differences (Pigot et al. 2020), but further studies could revisit latitudinal effects on beak evolution using methods that account for curvature, bill tip differences and other details of beak structure (Cooney *et al*. 2017a).

### Competition drives beak shape divergence across latitudes

Based on linear measurements, we report evidence that beak shape evolution is faster in sympatry than allopatry, suggesting a role for competition in driving divergence by ecological character displacement or ecological sorting. Importantly, we show that the relationship between beak shape divergence and sympatry is similar in the tropics and temperate zone. This is not consistent with the prediction of the biotic interaction hypothesis that competition is stronger in the tropics, though the possibility remains that there are latitudinal differences in diffuse competition (as opposed to the pairwise competition that we measured). While we also find that beak size divergence is greater in sympatric sister pairs than allopatric sister pairs, this is explained not by competition but by the older age of the sympatric sister pairs. Given that sympatric and allopatric sister pairs have similar rates of beak size evolution and relatively minor differences in beak shape evolution, our results suggest that character displacement (and ecological sorting) play a relatively minor role in global patterns of bird beak evolution (Tobias *et al*. 2020).

### Evolutionary rates are typically fastest in the temperate zone

Our study adds to a growing body of evidence documenting that recent evolutionary rates are often faster in the temperate zone than in the tropics. While early empirical tests reported faster molecular evolution in the tropics (e.g., Gillooly *et al*. 2005; Wright *et al*. 2006), a recent comprehensive analysis of ~8,000 taxon pairs from six phyla of animals found that latitudinal differences in the rate of molecular evolution are indeed faster in the tropics, but only just (faster in the tropics in 51.6% of comparisons vs. faster in the temperate zone in 48.4% of comparisons)(Orton *et al*. 2019). In contrast, trait evolution and recent speciation rates are faster in the temperate zone than the tropics in all published studies to date. All studies of trait evolution are on birds (Martin *et al*. 2010; Weir & Wheatcroft 2011; Lawson & Weir 2014; Weir & Price 2019); studies in additional taxonomic groups are necessary to test the generality of this pattern. However, recent speciation rates are fastest at high latitudes in angiosperms (Igea & Tanentzap 2020), marine fishes (Rabosky *et al*. 2018) and in both birds and mammals (Weir & Schluter 2007). This same pattern, replicated across different taxonomic groups, directly contradicts predictions arising from fast tropical evolution hypotheses, and instead is consistent with the idea that ecological opportunity has been greater on average at high latitudes in recent evolutionary history.

## Conclusion

Explaining the latitudinal diversity gradient is a longstanding goal of evolutionary ecology. The generality of the latitudinal diversity gradient across taxa, space and time suggests the tantalizing possibility of a common explanation. Since Darwin and Wallace, generations of biologists have added new hypotheses to a growing list of contenders (reviewed by Pianka 1966; Rohde 1992; Mittelbach *et al*. 2007). We argue that, on the strength of current evidence, hypotheses that invoke faster evolution in the tropics to explain the latitudinal diversity gradient should be rejected. Evolutionary hypotheses that invoke slower extinction in the tropics (rather than faster evolution) remain viable, though difficult to test. For example, a latitudinal gradient in the strength of biotic interactions in the tropics may help to explain how high tropical diversity is maintained (by lower extinction) rather than generated (by faster evolution). Future studies should consider how biotic interactions influence extinction rates in a latitudinal context.

## Acknowledgements

Comments from Jason Weir, the Schluter lab group and Ralf Yorque greatly improved this manuscript. BGF was supported by postdoctoral fellowships from the Biodiversity Research Centre and Banting Canada (#379958). Morphological data collection was conducted by Catherine Sheard, Monte Neate-Clegg, and Nico Alioravainen, among many others, and funded by the UK Natural Environment Research Council (NE/I028068/1 and NE/P004512/1 to JAT). None of our funders had any influence on the content of the submitted manuscript, and none of our funders required approval of the final manuscript to be published.

## Author contributions

JAT assembled morphological dataset, BGF and DS conducted analyses, BGF wrote the first draft of the manuscript and DS and JAT contributed substantially to revisions.

## Data accessibility statement

All data and R scripts will be uploaded to Dryad upon acceptance.

## Notes

### Competing Interest Statement

The authors have declared no competing interest.

